# Elasticity generates indissoluble biomolecular condensates

**DOI:** 10.1101/2022.02.09.479808

**Authors:** Lingyu Meng, Jie Lin

**Affiliations:** Peking-Tsinghua Center for Life Sciences, Peking University, Beijing, China; Center for Quantitative Biology, Peking University, Beijing, China

## Abstract

While biomolecular condensates are often liquid-like, many experiments found that condensates also exhibit solid-like behaviors, making them indissoluble in conditions liquid condensates dissolve. Despite the biological significance of indissoluble condensates to cellular fitness, the mechanisms underlying the indissolubility of solid-like condensates are still unclear. In this work, we study the effects of elasticity on the dissolution of biomolecular condensates. We demonstrate that the bulk stress inside condensates may prevent the condensates from dissolution and obtain a new mechanical equilibrium condition of elastic condensates. Moreover, we theoretically predict a phase diagram of indissolubility for biomolecular condensates and identify a minimum bulk modulus for the condensates to be indissoluble. To verify our theories, we simulate the two-fluid model in which the slow component corresponding to biomolecules generates elastic stress. Our theoretical predictions are nicely confirmed and independent of microscopic details. Our works show that elasticity makes biomolecular condensates less prone to dissolution.

---

Biomolecular condensates are ubiquitous in various organisms, usually composed of proteins and RNAs [1–9]. They often have crucial biological functions [8, 10–16], such as adaptive responses to stresses, accelerating biochemical reactions, and sequestering molecules from reactions. Therefore, the accurate regulation of biomolecular condensates’ formation and dissolution is critical. Meanwhile, experiments have also found that biomolecular condensates are viscoelastic [17]: they are solid-like on a short time scale and liquid-like on a long time scale. Interestingly, they often exhibit aging behaviors, and the viscoelastic relaxation time, which separates solid and liquid behaviors, increases over time [18]. Indeed, aged condensates may become indissoluble or infusible in conditions where newly formed condensates can easily dissolve or fuse [3, 4, 9, 10, 14, 19–21]. The resistance to dissolution of solid-like condensates is particularly significant when the condensates are induced by deep supersaturation [7]. Dissolution of solid-like condensates may need assistance by energy-consuming enzymes [20, 22–24], therefore, reducing cellular fitness. Moreover, failure to dissolve condensates during mitosis leads to aberrant condensates that cause cell-cycle arrest and ultimately cell death [25].

Despite the importance of solid-like nature on the dissolution of condensates [15, 18], theoretical studies on the formation and dissolution of biomolecular condensates have so far been limited to fluid models, in which the elastic nature of condensates are neglected [26]. Among the most common theoretical models, the hydrodynamic model involving diffusion and advection, often called Model H [26, 27], can successfully incorporate the physics needed to describe the dynamics of phase separation and droplet growth. However, it does not include the elastic nature of condensates. Therefore, it cannot explain the indissolubility of solid-like condensates in conditions where liquid condensates dissolve.

In this work, we investigate the effects of elasticity on the dissolubility of biomolecular condensates, combining both theories and numerical simulations. In the following, we first introduce our theoretical frameworks, focusing on elastic condensates subject to an abrupt control parameter change, e.g., some post-translational modifications that reduce the attractive strength between biomolecules. Without elasticity, they are supposed to dissolve. We derive the mechanical equilibrium conditions for elastic condensates and find that the bulk stress may prevent the dissolution and render the condensate indissoluble. To test our theoretical predictions, we simulate the two-fluid model [28, 29] beyond the traditional Model H, in which the biomolecule velocity field and the elastic stress are dynamically coupled. Numerical simulations of the two-fluid model nicely confirm our theories. Our theories’ validity is independent of the microscopic details, such as the free energy forms of the biomolecule density field, as we confirm numerically.

Furthermore, we theoretically predict a phase diagram of indissolubility as a function of effective temperature and the condensate’s bulk modulus. We demonstrate a minimum bulk modulus for the condensate to be indissoluble. Numerical simulations nicely confirm our predictions regarding the boundary between the dissoluble and indissoluble phases. Our results suggest that the dissolution of elastic condensates can be facilitated by decreasing biomolecules’ attractive strength and lowering the condensates’ bulk moduli. Finally, we discuss the biological implications of our work and propose potential experiments to test our predictions.

## Equilibrium conditions of elastic condensates

Biomolecular condensates usually have well-defined viscoelastic relaxation times, below which the condensates behave as elastic materials [17, 18]. In this work, we simplify the problem by considering aged condensates with their viscoelastic relaxation times much longer than the time scales of biological interests, e.g., the duration of cell-cycle phases. Therefore, the viscoelastic relaxation time can be taken as infinite, which is the main focus of this work. We introduce an abrupt reduction to the attractive strength between the biomolecules. Biologically, one of the most relevant perturbations is the post-translational modification such as phosphorylation, which introduces Coulomb repulsive interaction between biomolecular monomers and reduces the attractive interaction [5, 13, 30]. Other perturbations that can lead to similar effects include changing temperature, pH, and salt concentrations. In the absence of elasticity, the condensates will dissolve because the free energy of biomolecule density changes from a form with two minimums to a form with only one minimum (Figure 1). However, as shown in the following, the elastic force may prevent the dissolution.

**FIG. 1.**
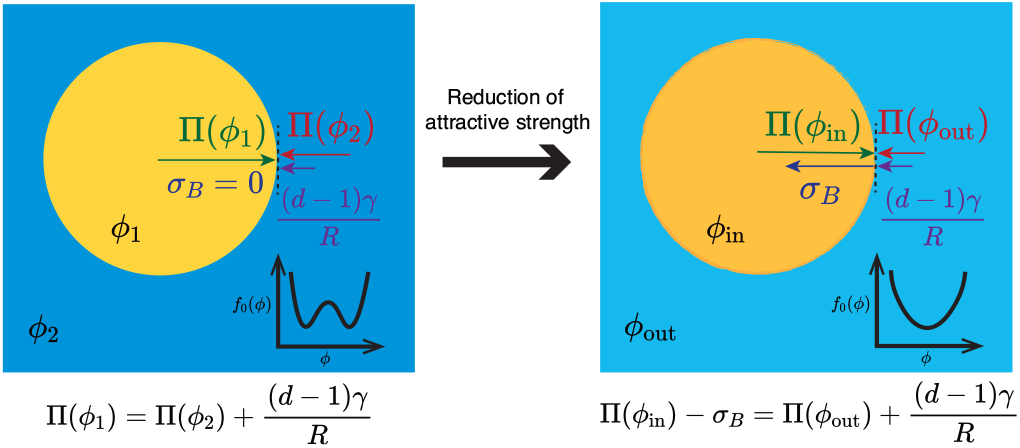
Mechanical equilibrium conditions of elastic condensates. Liquid-like condensates are initially formed, which then become aged and solid-like. An abrupt change in the attractive strength between biomolecules is introduced, corresponding to a change in the shape of the biomolecule free energy *f*_0_(*ϕ*) from two minimums to a single minimum. Here *ϕ* is the biomolecule density. The condensate’s bulk stress *σ_B_* is involved in the new equilibrium condition.

In liquid-liquid phase separation, a stable condensate requires mechanical equilibrium [31, 32], such that Π_in_ = Π_out_ + (*d* – 1)*γ*/*R*. Here, Π_in_ (_out_) is the osmotic pressure inside (outside) the condensate, *d* is the spatial dimension, *γ* is the surface tension constant and *R* is the condensate radius. In the presence of elastic stress, we need to take account of the elastic energy *F*_el_ the condensate so that the equilibrium condition becomes

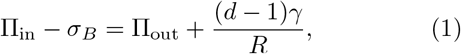

Here *σ_B_* = ∂*F*_el_(*V*_in_)/∂*V*_in_ is the bulk stress inside the condensate due to elasticity where *V*_in_ is the volume of the condensate (Figure 1). As we show later, the inclusion of bulk stress compensates the imbalance of osmotic pressures. To find the expression of *σ_B_*, we use the constitutive equation of the bulk stress and the continuity equation of biomolecule density (*ϕ*)

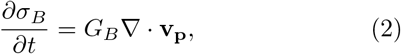

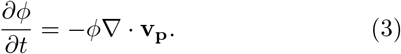

Here **v_p_** is the velocity field of the biomolecules, which is responsible for the bulk stress. *G_B_* is the bulk modulus. In writing the above two equations, we assume that *ϕ* and *σ_B_* are uniform inside the condensate, which we confirm numerically later. Combining Eqs. (2, 3), we obtain

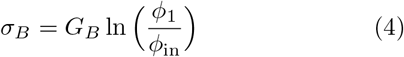

where *ϕ*_1_ and *ϕ*_in_ are the biomolecule densities inside condensates before and after the reduction of attractive strength between the biomolecules. The equilibrium condition must be satisfied for all condensates in a system of multiple condensates with the bulk stress determined by Eq. (4), and *Π*_in_ and *Π*_out_ determined by *ϕ*_in_ and *ϕ*_out_, respectively. Each condensate can have different *ϕ*_in_’s although they share the same biomolecule density *ϕ*_out_ outside them. Given the radii R’s and the densities *ϕ*_in_’s of each condensate, *ϕ*_out_ can be calcu-lated using the conservation of total molecular number: 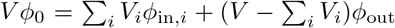. Here, *V* is the total volume, *V_i_* is the volume of condensate *i* assuming a spherical shape, and the summation is over all condensates. *ϕ*_0_ is the average density over the total volume.

To uniquely determine the densities inside condensates, we still need one more equation. For liquid condensates, it is a uniform chemical potential. However, in our case, the condensate is solid, so the exchange of molecules is suppressed as soon as the mechanical equilibrium is established [3, 8, 15]. Instead, we propose that condensate size does not change upon the weakening of attractive interaction between biomolecules, namely, *R* = *R*_0_, where *R* and *R*_0_ are the radii of an elastic condensate before and after the condition changes. We confirm this assumption numerically later. We note that the condensate’s constant radius and decreased density do not conflict with the suppressed exchange of molecules since the density change happens when the system is initially out-of-equilibrium after the attractive strength suddenly decreases. As a result, the density decreases initially until the condensate reaches mechanical equilibrium. Finally, we remark that in our case, the bulk stress inside the condensate stabilizes the condensate, in contrast to the bulk stress outside a condensate, e.g., due to the surrounding polymer network that suppresses the formation of condensates [33–40].

## Simulations of the two-fluid model

We numerically simulate the two-fluid model in two dimensions [28, 29] to test our theories with two components: the slow component corresponding to the biomolecules and the fast component corresponding to the solvent. It is the biomolecule component that generates the elastic stress. The average velocity field **v** = *ϕ***v_p_** + (1 – *ϕ*)**v_s_** where **v_p_** are **v_s_** are respectively the biomolecule and solvent velocity field. In the two-fluid model, the biomolecule density field *ϕ* is spatially dependent with its value between 0 and 1. The dynamics of the biomolecule density and velocity field follows (see details in Supplementary Information)

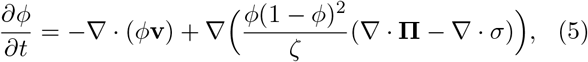

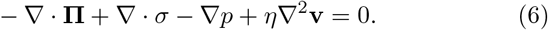

Here, *ζ* is the friction constant between biomolecules and solvent, and *η* is the viscosity. The pressure *p* is determined by the incompressible condition: ∇ · **v** = 0. The stress tensor *σ* = *σ_S_*+*σ_B_***I** where *σ_B_* is the bulk stress and *σ_S_* is the shear stress tensor. They follow the Maxwellian dynamics such that (Supplementary Information),

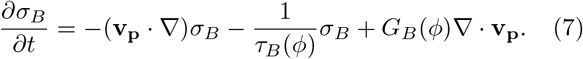

where *G_B_* is the bulk modulus. To simulate elastic condensates, we take the relaxation time of bulk stress to be diverging at a critical density *ϕ*_c_ such that 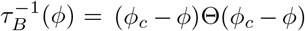, where Θ(*x*) is the Heaviside function. We remark that if we take the stress tensor to be zero in Eqs. (5, 6), they are reduced to the classical Model H [27]. The osmotic stress tensor is determined by the biomolecule free energy 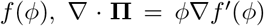 where 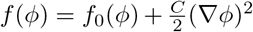 and *C* is a constant. If not mentioned explicitly, we use the Flory-Huggins free energy density: 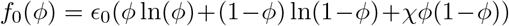. Here *ϵ*_0_ = *k_B_T/V*_0_ where *k_B_* is the Boltzmann constant, *T* is the temperature, and *V*_0_ is the biomolecular monomer volume. We note that the condition of stable liquid condensate for the Flory-Huggins model is that the parameter *χ* > 2. We use the osmotic pressure *Π* to represent the scalar osmotic stress computed from *f*_0_ (*ϕ*), 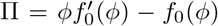. By non-dimensionalizing Eqs. (5, 6), we choose the unit of elastic modulus as *ϵ*_0_, the time unit *t*_0_ = *η*/*ϵ*_0_, and the length unit 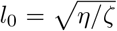 (see estimations in Supplementary Information).

We note that the osmotic stress and the elastic stress are fundamentally different as the osmotic stress is a function of the instantaneous density field while the elastic stress depends on the accumulated change of the density field. One may attempt to compute an effective elastic free energy that is a function of *ϕ* so that the elastic bulk stress is included in the osmotic pressure as 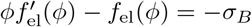. However, the resulting effective chemical potential 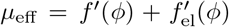 is not uniform across the system, which suggests that the elastic stress cannot be included in the free energy as a function of the density field (Figure S1).

## Tests of theoretical predictions

We simulate multiple coexisting condensates, first generated by the two-fluid model without elasticity from a uniform density *ϕ*_0_ as the initial condition (see details in Supplementary Information). The average density is therefore set by the initial condition. The initial Flory-Huggins parameter *χ_i_* = 3. We add elasticity to the condensates after the formation of multiple spherical condensates to mimic the aging process. After a short time, we change *χ* to *χ_f_* = 1.5 so that the free energy *f*_0_(*ϕ*) changes from a form with two minimums to a form with only one minimum (Figure 1).

After the reduction of *χ*, we find that these condensates are indissoluble under the parameters we take. The density field inside condensates is indeed uniform as assumed (Figure 2a). We also confirm our assumptions of uniform bulk stress (Figure 2c) and constant radii (Figure S2). An example of simulations is shown in Movie S1. The osmotic pressure is significantly different across the boundaries of condensates (Figure 2b). For liquid condensates, they will quickly dissolve due to the large pressure difference. In contrast, the bulk stress balances the osmotic pressure difference for elastic condensates. Indeed, we find that the difference between the osmotic pressure and bulk stress (Π – *σ_B_*) is uniform across the boundaries (Figure 2d). We note that the uniform Π–*σ_B_* is not valid near the condensate boundaries due to the surface tension. Using the variable sizes of condensates, we compute the radius dependence of surface tension constant *γ*, and find that *γ* converges to a constant value in the large radius limit (Figure 3a), suggesting that it is well defined in the thermodynamic limit.

**FIG. 2.**
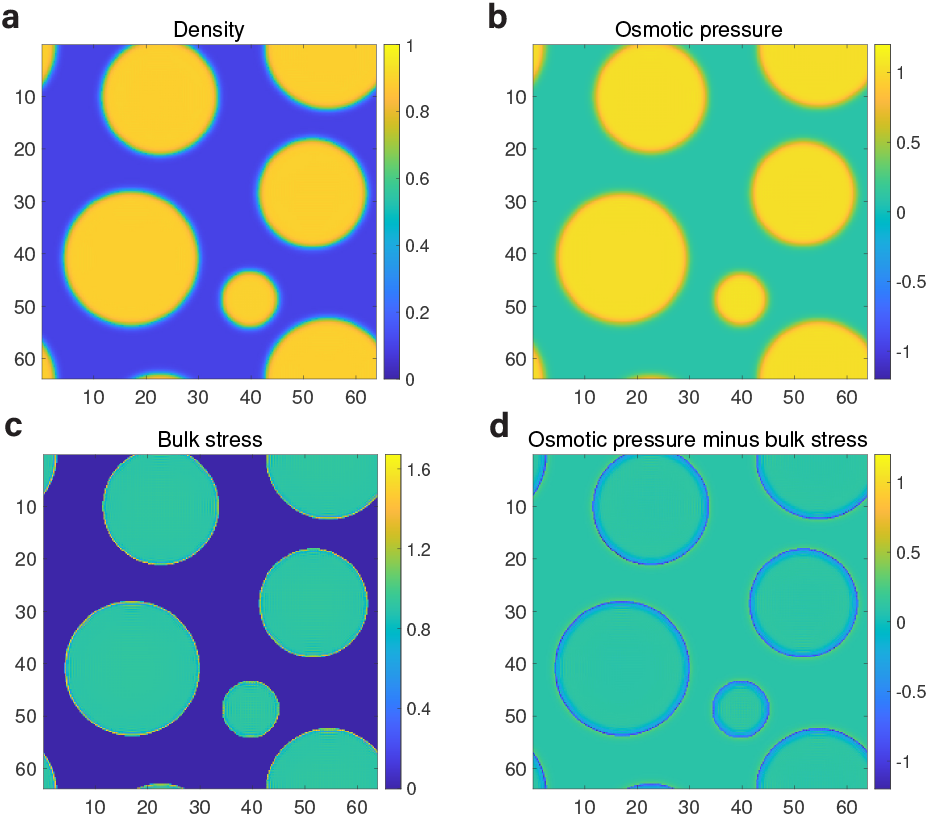
Simulations of multiple coexisting elastic condensates. (a) The density field *ϕ* after decreasing the *χ* parameter in the Flory-Huggins free energy from 3.0 to 1.5. For liquid condensates, they will simply dissolve. In contrast, the elastic condensates can be indissoluble. (b) The osmotic pressure *ϕ* from the same simulation of (a). (c) The bulk stress *σ_B_* from the same simulation of (a). (d) *ϕ* – *σ_B_* from the same simulation of (a). In this figure, we take *ϕ*_0_ = 0.45, *G_B_* = 20, *G_S_* = 20, and *ϕ_c_* = 0.5.

**FIG. 3.**
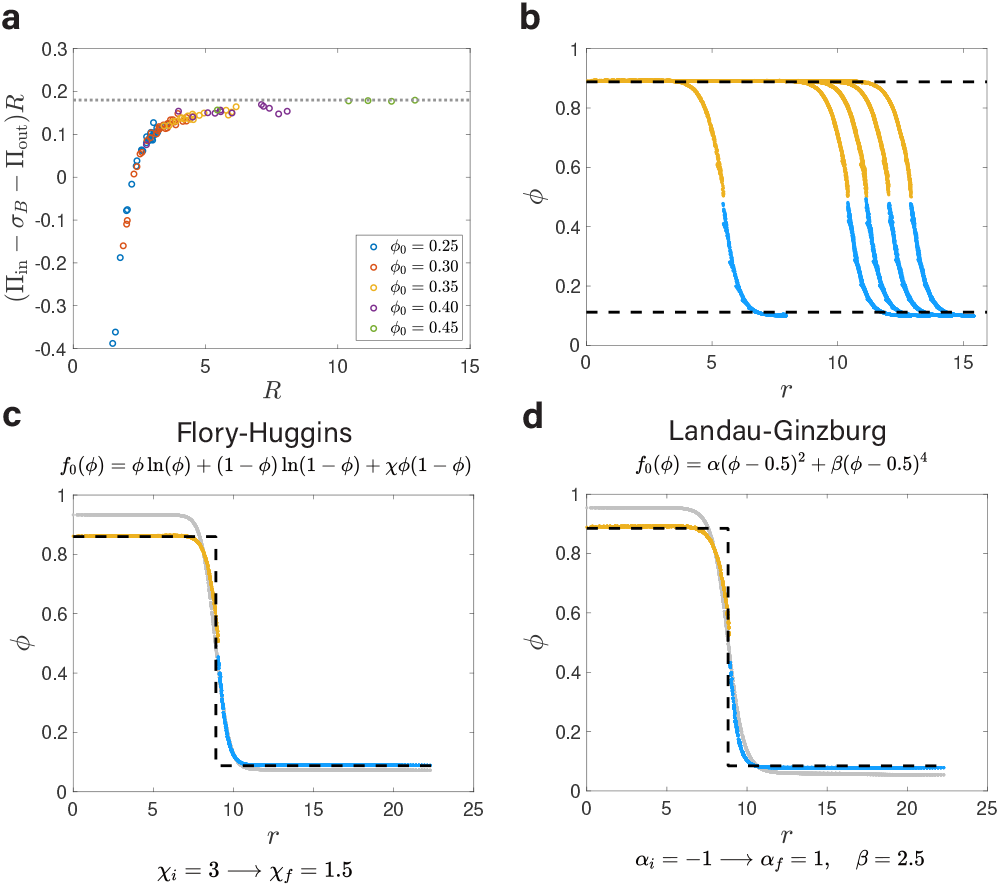
Computations of the surface tension constant *γ* and predictions of the density field using different types of free energy. (a) The inferred surface tension constant *γ* = (Π_in_ – *σ_B_* – Π_out_)*R* approaches an asymptotic value in the large radius limit. (b) A comparison of the theoretical predictions of *ϕ*_in_ and *ϕ*_out_ (black dashed lines) and the simulations (yellow dots above *ϕ_c_* and blue dots below *ϕ_c_*) [same as Figure 2(a)]. Each yellow and blue curve represents one condensate and *r* is the distance from the condensate center. (c) For the Flory-Huggins free energy, the initial density field (gray dots) cannot be maintained after *χ* decreases and the final density field is established. (d) For the Landau-Ginzburg free energy, the equilibrium density field can also be predicted by our theories after the control parameter *α* increases from –1 to 1. In (a) and (b), *G_B_* = 20, *χ_i_* = 3.0, *χ_f_* = 1.5. *ϕ*_0_ = 0.45 in (b). In (c) and (d), we simulate a single condensate (see details in Supplementary Information) with *R*_0_ = 9. *G_B_* = 10 in (c) and *G_B_* = 20 in (d). In all figures, *G_S_* = 20 and *ϕ_c_* = 0.5.

To compute the predicted densities inside and outside condensates, in principle, we need to solve *n* equations of Eq. (1) with a shared Π_out_(*ϕ*_out_) because each condensate has a different radius. Here, *n* is the number of condensates. However, if we neglect the contribution of surface tension in Eq. (1), we can combine all condensates to find the common *ϕ*_in_ that works for all condensates. We find that the surface tension constant is relatively small in our simulations, so our predictions of the densities *ϕ_in_* and *ϕ*_out_ with *γ* = 0 (Figure 3b) are very close to the predictions with a finite *γ* (Figure S3). Therefore, we neglect the surface tension in the following theoretical calculations.

To test the generality of our theories, we also use the Landau-Ginzburg free energy and find that our theories are equally applicable (Figure 3c, d). In both cases, the condensates can be indissoluble after changing the forms of free energies from two minimums to a shape with only one minimum. We find that the numerical densities *ϕ* as a function of the distance from the condensate center precisely match the theoretical predictions. We also test our theories using asymmetric Flory-Huggins model and again obtain satisfying agreements between simulations and theoretical predictions (Figure S4a).

To test the effects of the shear modulus, we repeat the above simulations with the Flory-Huggins free energy and change the shear modulus with the same bulk modulus. We find that the density distributions are insensitive to the values of shear moduli (Figure S5), corroborating the major role of bulk stress in the mechanical equilibrium conditions. Finally, we also test the effects of the critical density and find that our results are valid independent of *ϕ_c_* (Figure S4b, S6-S8).

## Phase diagram of indissolubility

In the following, we systematically investigate the indissoluble conditions of elastic condensates. We compute the theoretically predicted *ϕ*_in_ after *χ* is lowered from *χ_i_* = 3 to *χ_f_* as a function of *G_B_* using Eq. (1). We find that *ϕ*_in_ decreases as *G_B_* decreases (Figure S9a). From this calculation, we predict the critical bulk modulus *G_B, c_* for the condensate to be indissoluble given *χ_f_*, based on the condition *ϕ*_in_ > *ϕ_c_* where *ϕ_c_* is the critical density above which the condensates become elastic. Because both *χ* and *G_B_* appear linearly in Eq. (1), the phase boundary separating the dissoluble and indissoluble phase must be linear in the *χ_f_*-*G_B_* parameter space.

To test our predictions, we simulate a single condensate with two control parameters *χ_f_* and *G_B_* and monitor its dissolution dynamics after *χ* is reduced from *χ_i_* = 3 to *χ_f_*. We label the condensate as dissoluble or indissoluble depending on if the system becomes uniform or not after a long waiting time *t* = 10^4^ (see examples in Figure S9b, c and Movie S2). As expected, when *G_B_* = 0, the condensate is stable only if *χ_f_* > 2. For *χ_f_* < 2, the condensate becomes indissoluble if the bulk modulus is larger than a critical value. The numerically simulated phase diagram nicely matches the predicted phase diagram (Figure 4a). Our results are not sensitive to the values of *ϕ_c_* as we get similar results using different *ϕ_c_* (Figure S10).

**FIG. 4.**
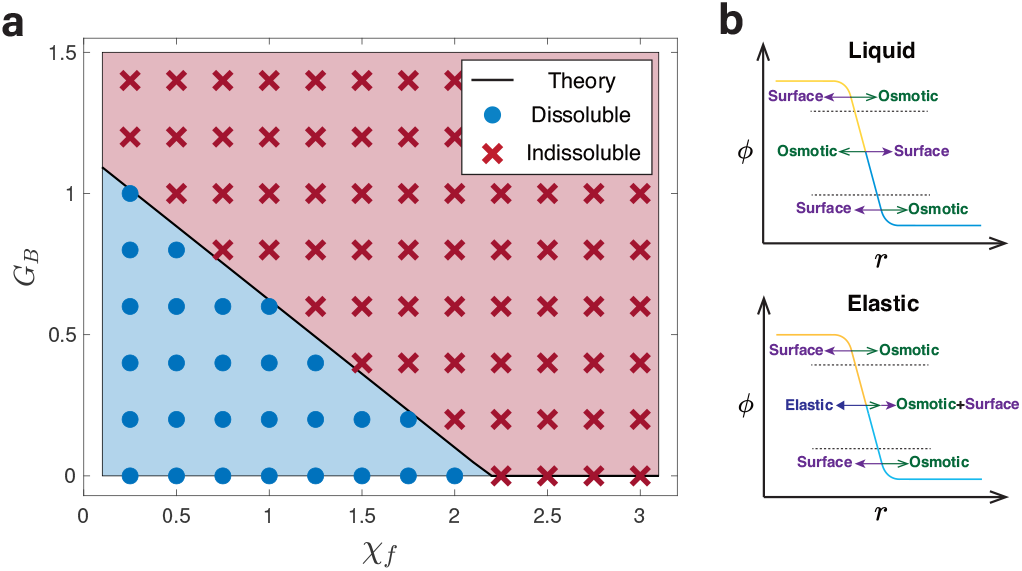
A critical bulk modulus *G_B,c_* above which condensates are indissoluble. (a) Phase diagram of condensate indissolubility with control parameters *χ_f_* and *G_B_*. The theoretical predicted *G_B,c_* is the black line and the simulation results are the blue dots and red crosses. (b) Schematics for the force balance in the crossover regime of condensates.

We find that to match the predicted phase boundary to the simulations accurately, we need to solve the equation *ϕ*_in_ = *θϕ_c_* with *θ* = 1.1 to find the critical bulk modulus (Figure 4a and Figure S9a). To understand why θ is close but slightly larger than 1, we remark that for the elastic condensate to be stable, *ϕ*_in_ must be larger than *ϕ_c_* to ensure force balance across the condensate boundary. The biomolecules are subject to two types of force: the force from the gradient of the osmotic tensor (-∇ · **∏**) and the force from the gradient of the elastic stress (∇ · *σ*). We can further decompose the former force into two parts, one is from the free energy *f*_0_(*ϕ*), which we call the osmotic force, and the other is from the 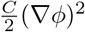 term in the free energy, which we call the surface tension force. For a liquid condensate, the osmotic force always balances the surface tension force across the condensate boundary. Because the osmotic pressure is a non-monotonic function of density, in this case, the crossover regime can be separated into three parts in which both the surface tension force and the osmotic force change their signs (see the schematic in Figure 4b and numerical simulations in Figure S11a). For an elastic condensate, the osmotic force always points outwards from the condensates since the osmotic pressure is now a monotonically increasing function of *ϕ*, while the surface tension force still changes its sign. Therefore, in this case, an inward elastic force must exist to balance the sum of surface tension force and osmotic force in the crossover regime (Figure 4b and Figure S11b). In conclusion, *ϕ*_in_ should be larger than *ϕ_c_* to ensure a finite elastic force in the crossover regime; therefore, *θ* ≳ 1.

## Discussion

Our work provides the first mechanistic understanding of the indissolubility of solid-like biomolecular condensates. We show that the bulk stress can balance the osmotic pressure difference inside and outside the condensates, therefore preventing the dissolution. Numerical simulations of the two-fluid model nicely confirm our theoretical predictions of the mechanical equilibrium condition. Moreover, we theoretically and numerically obtain a phase diagram of indissolubility for elastic condensates and obtain a minimum bulk modulus for condensates to be indissoluble. There remain some open questions, including the effects of shear modulus. While we numerically find that our simulations are mostly independent of the values of shear moduli, a small but finite shear modulus is nevertheless necessary to maintain the spherical shapes of condensates.

Finally, our results demonstrate that the bulk modulus is the primary material property determining condensates’ indissolubility. The phase diagram of indissolubility (Figure 4) can be experimentally tested, e.g., by measuring the critical pH or salt concentrations (corresponding to the *χ* parameter in the Flory-Huggins free energy) to dissolve the condensates and in the meantime, measuring the bulk modulus separately. We note that the shear modulus of gel is usually the order of *nk_B_ T* where *n* is the cross-linker density, and meanwhile, the bulk modulus is typically hundreds of shear modulus [31]. Note that the unit of the bulk modulus is *k_B_T/V*_0_ in the two-fluid model, which suggests that the critical bulk moduli that are of order 1 in the phase diagram (Figure 4d) may be biologically relevant. Our results may have implications for developing condensate-targeting drugs to change condensate properties inside cells, e.g., lowering the bulk moduli to dissolve irreversible condensates.

We thank Xiangyu Kuang, Sheng Mao, Yiyang Ye, and Lei Zhang for useful discussions related to this work. The research was supported by grants from Peking-Tsinghua Center for Life Sciences.

## References

[1] C. P. Brangwynne, C. R. Eckmann, D. S. Courson, A. Rybarska, C. Hoege, J. Gharakhani, F. Jülicher, and A. A. Hyman, Science 324, 1729 (2009).

[2] C. P. Brangwynne, P. Tompa, and R. V. Pappu, Nature Physics 11, 899 (2015).

[3] A. Patel, H. O. Lee, L. Jawerth, S. Maharana, M. Jahnel, M. Y. Hein, S. Stoynov, J. Mahamid, S. Saha, T. M. Franzmann, et al., Cell 162, 1066 (2015).

[4] Y. Lin, D. S. Protter, M. K. Rosen, and R. Parker, Molecular cell 60, 208 (2015).

[5] S. F. Banani, H. O. Lee, A. A. Hyman, and M. K. Rosen, Nature reviews Molecular cell biology 18, 285 (2017).

[6] J. B. Woodruff, B. F. Gomes, P. O. Widlund, J. Mahamid, A. Honigmann, and A. A. Hyman, Cell 169, 1066 (2017).

[7] Y. Shin, J. Berry, N. Pannucci, M. P. Haataja, J. E. Toettcher, and C. P. Brangwynne, Cell 168, 159 (2017).

[8] T. M. Franzmann, M. Jahnel, A. Pozniakovsky, J. Mahamid, A. S. Holehouse, E. Nüske, D. Richter, W. Baumeister, S. W. Grill, R. V. Pappu, A. A. Hyman, and S. Alberti, Science 359(2018).

[9] J. Wang, J.-M. Choi, A. S. Holehouse, H. O. Lee, X. Zhang, M. Jahnel, S. Maharana, R. Lemaitre, A. Pozniakovsky, D. Drechsel, et al., Cell 174, 688 (2018).

[10] S. Alberti and A. A. Hyman, BioEssays 38, 959 (2016).

[11] Y. Shin and C. P. Brangwynne, Science 357(2017).

[12] S. Alberti and D. Dormann, Annual Review of Genetics 53, 171 (2019), pMID: 31430179.

[13] S. Alberti, A. Gladfelter, and T. Mittag, Cell 176, 419 (2019).

[14] C. Iserman, C. Desroches Altamirano, C. Jegers, U. Friedrich, T. Zarin, A. W. Fritsch, M. Mittasch, A. Domingues, L. Hersemann, M. Jahnel, D. Richter, U.-P. Guenther, M. W. Hentze, A. M. Moses, A. A. Hyman, G. Kramer, M. Kreysing, T. M. Franzmann, and S. Alberti, Cell 181, 818 (2020).

[15] S. Alberti and A. A. Hyman, Nature Reviews Molecular Cell Biology 22, 196 (2021).

[16] A. S. Lyon, W. B. Peeples, and M. K. Rosen, Nature Reviews Molecular Cell Biology 22, 215 (2021).

[17] L. M. Jawerth, M. Ijavi, M. Ruer, S. Saha, M. Jahnel, A. A. Hyman, F. Jülicher, and E. Fischer-Friedrich, Phys. Rev. Lett. 121, 258101 (2018).

[18] L. Jawerth, E. Fischer-Friedrich, S. Saha, J. Wang, T. Franzmann, X. Zhang, J. Sachweh, M. Ruer, M. Ijavi, S. Saha, J. Mahamid, A. A. Hyman, and F. Julicher, Science 370, 1317 (2020).

[19] A. Molliex, J. Temirov, J. Lee, M. Coughlin, A. P. Kana-garaj, H. J. Kim, T. Mittag, and J. P. Taylor, Cell 163, 123 (2015).

[20] J. A. Riback, C. D. Katanski, J. L. Kear-Scott, E. V. Pilipenko, A. E. Rojek, T. R. Sosnick, and D. A. Drummond, Cell 168, 1028 (2017).

[21] T. S. Harmon, A. S. Holehouse, M. K. Rosen, and R. V. Pappu, eLife 6, e30294 (2017).

[22] S. Jain, J. R. Wheeler, R. W. Walters, A. Agrawal, A. Barsic, and R. Parker, Cell 164, 487 (2016).

[23] M. Hondele, R. Sachdev, S. Heinrich, J. Wang, P. Vallotton, B. M. Fontoura, and K. Weis, Nature 573, 144 (2019).

[24] D. Tauber, G. Tauber, A. Khong, B. Van Treeck, J. Pelletier, and R. Parker, Cell 180, 411 (2020).

[25] A. K. Rai, J.-X. Chen, M. Selbach, and L. Pelkmans, Nature 559, 211 (2018).

[26] J. Berry, C. P. Brangwynne, and M. Haataja, Reports on Progress in Physics 81, 046601 (2018).

[27] P. C. Hohenberg and B. I. Halperin, Rev. Mod. Phys. 49, 435 (1977).

[28] H. Tanaka, Journal of Physics: Condensed Matter 12, R207 (2000).

[29] H. Tanaka and T. Araki, Chemical Engineering Science 61, 2108 (2006).

[30] J. Söding, D. Zwicker, S. Sohrabi-Jahromi, M. Boehning, and J. Kirschbaum, Trends in Cell Biology 30, 4 (2020).

[31] M. Doi, Soft matter physics (Oxford University Press, 2013).

[32] A. W. Fritsch, A. F. Diaz-Delgadillo, O. Adame-Arana, C. Hoege, M. Mittasch, M. Kreysing, M. Leaver, A. A. Hyman, F. Jülicher, and C. A. Weber, Proceedings of the National Academy of Sciences 118(2021).

[33] R. W. Style, T. Sai, N. Fanelli, M. Ijavi, K. Smith-Mannschott, Q. Xu, L. A. Wilen, and E. R. Dufresne, Phys. Rev. X 8, 011028 (2018).

[34] K. A. Rosowski, E. Vidal-Henriquez, D. Zwicker, R. W. Style, and E. R. Dufresne, Soft Matter 16, 5892 (2020).

[35] X. Wei, J. Zhou, Y. Wang, and F. Meng, Phys. Rev. Lett. 125, 268001 (2020).

[36] Y. Zhang, D. S. W. Lee, Y. Meir, C. P. Brangwynne, and N. S. Wingreen, Phys. Rev. Lett. 126, 258102 (2021).

[37] S. Biswas, B. Mukherjee, and B. Chakrabarti, arXiv preprint arXiv:2104.00651 (2021).

[38] E. Vidal-Henriquez and D. Zwicker, Proceedings of the National Academy of Sciences 118, e2102014118 (2021).

[39] P. Ronceray, S. Mao, A. Košmrlj, and M. P. Haataja, Europhysics Letters 137, 67001 (2022).

[40] M. Kothari and T. Cohen, arXiv preprint arXiv:2201.04105 (2022).

