## Supplementary Information for "Elasticity generates indissoluble biomolecular condensates"

(Dated: July 11, 2022)

### A. Details of the two-fluid model

Numerical simulations of viscoelastic phase separation are based on the two-fluid model [1, 2], which considers the dynamics of biomolecule velocity $\mathbf{v_p}$, solvent velocity $\mathbf{v_s}$ and the average velocity $\mathbf{v} = \phi\mathbf{v_p} + (1-\phi)\mathbf{v_s}$. This model is derived by minimizing the Rayleighian $R$ [1–3], the sum of the energy dissipation function $\Phi$ and the temporal changing rate of the free energy $\dot{F}$. The energy dissipation function $\Phi$ consists of two parts, which are respectively the friction between biomolecules and solvent $\Phi_1$, and the overall viscous dissipation of the solution $\Phi_2$

$$\Phi_1 = \int \mathrm{d}\mathbf{r}\, \frac{\zeta}{2}(\mathbf{v_p} - \mathbf{v_s})^2 = \int \mathrm{d}\mathbf{r}\, \frac{1}{2}\zeta \frac{(\mathbf{v_p} - \mathbf{v})^2}{(1-\phi)^2}, \tag{S1}$$

$$\Phi_2 = \int \mathrm{d}\mathbf{r}\, \frac{\eta}{4}\big(\nabla\mathbf{v} + (\nabla\mathbf{v})^T\big) : \big(\nabla\mathbf{v} + (\nabla\mathbf{v})^T\big). \tag{S2}$$

The temporal changing rate of the mixing free energy $\dot{F}_{\text{mix}}$ is calculated through

$$\begin{aligned}\dot{F}_{\text{mix}} &= \int \mathrm{d}\mathbf{r}\, \dot{\phi} f'(\phi) = \int \mathrm{d}\mathbf{r}\, [-\nabla\cdot(\phi\mathbf{v_p})] f'(\phi) \\ &= \int \mathrm{d}\mathbf{r}\, [\phi\nabla f'(\phi))]\cdot\mathbf{v_p} = \int \mathrm{d}\mathbf{r}\, (\nabla\cdot\mathbf{\Pi})\cdot\mathbf{v_p},\end{aligned} \tag{S3}$$

involving the continuity equation of the density: $\dot{\phi} = -\nabla\cdot(\phi\mathbf{v_p})$. Here $f(\phi) = f_0(\phi) + \frac{C}{2}(\nabla\phi)^2$, where the unit of $f_0$ is $k_BT/V_0$ and $C$ is a constant. The elastic energy comes from the biomolecules, so its temporal changing rate is

$$\dot{F}_{\text{el}} = \int \mathrm{d}\mathbf{r}\, \sigma_{ij}\partial_j v_{p_i} = \int \mathrm{d}\mathbf{r}\, (-\nabla\cdot\sigma)\cdot\mathbf{v_p}. \tag{S4}$$

Combined with the constrain from the incompressible condition

$$\nabla\cdot\mathbf{v} = 0, \tag{S5}$$

and all the components mentioned above, the Rayleighian becomes:

$$\begin{aligned}R = \int \mathrm{d}\mathbf{r}\, \Big[ &-p(\nabla\cdot\mathbf{v}) + \frac{\zeta}{2}\frac{(\mathbf{v_p}-\mathbf{v})^2}{(1-\phi)^2} \\ &+ \frac{\eta}{4}\big(\nabla\mathbf{v} + (\nabla\mathbf{v})^T\big) : \big(\nabla\mathbf{v} + (\nabla\mathbf{v})^T\big) \\ &+ (\nabla\cdot\mathbf{\Pi})\cdot\mathbf{v_p} - (\nabla\cdot\sigma)\cdot\mathbf{v_p}\Big].\end{aligned} \tag{S6}$$

By setting the functional derivative of $R$ with $\mathbf{v_p}$ and $\mathbf{v}$ to be 0, we obtain the following equations:

$$\frac{\zeta}{(1-\phi)^2}(\mathbf{v_p} - \mathbf{v}) + \nabla\cdot\mathbf{\Pi} - \nabla\cdot\sigma = 0, \tag{S7}$$

$$\nabla p - \frac{\zeta}{(1-\phi)^2}(\mathbf{v_p} - \mathbf{v}) - \eta\nabla^2\mathbf{v} = 0. \tag{S8}$$

Finally, we rewrite the above equations and obtain

$$\frac{\partial\phi}{\partial t} = -\nabla\cdot(\phi\mathbf{v_p}), \tag{S9}$$

$$\mathbf{v_p} - \mathbf{v} = -\frac{(1-\phi)^2}{\zeta}(\nabla\cdot\mathbf{\Pi} - \nabla\cdot\sigma), \tag{S10}$$

$$-\nabla\cdot\mathbf{\Pi} + \nabla\cdot\sigma - \nabla p + \eta\nabla^2\mathbf{v} = 0, \tag{S11}$$

Clearly, Eq. (5) is obtained from Eq. (S9) and Eq. (S10). Combined with the incompressible condition $\nabla \cdot \mathbf{v} = 0$ and Eq. (S11), the average velocity $\mathbf{v}$ is calculated as

$$\mathbf{v}(\mathbf{r}) = \int \mathrm{d}\mathbf{r}' \, \mathbf{T}(\mathbf{r}-\mathbf{r}') \cdot (-\nabla \cdot \mathbf{\Pi}(\mathbf{r}') + \nabla \cdot \sigma(\mathbf{r}')), \tag{S12}$$

while $\mathbf{T}(\mathbf{k}) = \frac{1}{\eta|\mathbf{k}|^2}(\mathbf{I} - \frac{\mathbf{kk}}{|\mathbf{k}|^2})$ is the Oseen tensor in the Fourier space. We can then obtain the biomolecule velocity $\mathbf{v_p}$ with Eq. (S10). Therefore, the density $\phi$ in simulation can be updated by calculating $\mathbf{v_p}$ when the osmotic pressure $\mathbf{\Pi}$ and the stress $\sigma$ are known.

The shear stress tensor $\sigma_S$ also obeys the Maxwellian dynamics [2]:

$$\begin{aligned}\frac{\partial \sigma_S}{\partial t} =& -(\mathbf{v_p} \cdot \nabla)\sigma_S + \sigma_S \cdot \nabla \mathbf{v_p} + (\nabla \mathbf{v_p})^T \cdot \sigma_S \\ &- \frac{1}{\tau_S(\phi)}\sigma_S + G_S(\phi)(\nabla \mathbf{v_p} + (\nabla \mathbf{v_p})^T).\end{aligned} \tag{S13}$$

Here we make $\sigma_S$ traceless by setting $\sigma_S = \sigma_S - \mathrm{Tr}(\sigma_S)/d$. In our simulations, we first do not include elastic stress to form condensates by taking $G_S = G_B = 0$. We then introduce the elastic stress by taking

$$G_S(\phi) = G_S\phi^2, \tag{S14}$$

$$\tau_S^{-1}(\phi) = (\phi_c - \phi)\Theta(\phi_c - \phi), \tag{S15}$$

$$G_B(\phi) = G_B\Theta(\phi - \phi_c), \tag{S16}$$

$$\tau_B^{-1}(\phi) = (\phi_c - \phi)\Theta(\phi_c - \phi). \tag{S17}$$

Here $G_B$ and $G_S$ are constants. We assume a critical density $\phi_c$ above which the biomolecule network is percolated and becomes fully elastic with a finite bulk modulus and a diverging relaxation time.

We non-dimensionalize our model with the unit of elastic modulus as $\epsilon_0 = k_BT/V_0$, the time unit as $t_0 = \eta/\epsilon_0$ and the length unit as $l_0 = \sqrt{\eta/\zeta}$. If we estimate the biomolecule monomer volume as $V_0 = 1\mathrm{nm}^3$ and $T = 300\mathrm{K}$, we get $\epsilon_0 = 4.1 \times 10^6\mathrm{Pa}$. Taking the solvent viscosity as $\eta = 1.37 \times 10^{-2}\mathrm{Pa}\cdot\mathrm{s}$ [4], we obtain $t_0 = 3.3\mathrm{ns}$. While we do not have a good estimation of the friction constant $\zeta$, we find that if $l_0 = 10\mathrm{nm}$, $\zeta \approx 0.14\mathrm{pN}\cdot\mathrm{ns/nm}^4$, which appears a reasonable value for biomolecules.

### B. Details of numerical simulations

We perform numerical simulations in a 2D grid by solving the two-fluid model using the explicit Euler method with the periodic boundary condition on MATLAB. Simulations for a single condensate are in a $127 \times 127$ grid, and simulations for multiple condensates are in a $255 \times 255$ grid. In our simulations, we take $C/\epsilon_0 l_0^2 = 1$ and the grid size is $\Delta l = 0.25$. The time interval for the simulation is $\Delta t = 0.001$. The elastic stress is introduced at $t = 10^3$ for a single condensate and $t = 4 \times 10^3$ for multiple condensates. The control parameter $\chi$ is changed 50 time units after adding the elasticity. Eq. (S12) is solved with fast Fourier transformation, and other equations are calculated in real space. For the simulation of a single condensate, the initial density is set as $\phi_1$ when $r < R_0$ and as $\phi_2$ when $r > R_0$, where $r$ is the distance from the grid center, while $\phi_1$ and $\phi_2$ are the equilibrium densities of the free energy $f_0$ under $\chi_i$. The condensate has a sharp boundary initially and will soon evolve to the equilibrium density field with a continuous boundary. For the simulation of multiple condensates, we initially add a Gaussian noise with variance 0.001 to the uniform density field. In Figure 4a, simulations are initiated with a single condensate with $R_0 = 9$. The condensate is considered dissoluble if the variance of the $\phi$ field at $t = 10^4$ is less than 0.01.

In the theoretical predictions, $R_0$ is obtained as the radius of the region with $\phi > \phi_c$. When calculating $\sigma_B$ in Eq. (4), the rich-phase density of the free energy $f_0$ under $\chi_i$ is used as $\phi_1$, which is very close to the equilibrium density in the simulation and will not alter the theoretical predictions.


[1] H. Tanaka, Viscoelastic phase separation, Journal of Physics: Condensed Matter **12**, R207 (2000).
[2] H. Tanaka and T. Araki, Viscoelastic phase separation in soft matter: Numerical-simulation study on its physical mechanism, Chemical Engineering Science **61**, 2108 (2006).
[3] M. Doi, *Soft matter physics* (Oxford University Press, 2013).
[4] B. Parry, I. Surovtsev, M. Cabeen, C. O'Hern, E. Dufresne, and C. Jacobs-Wagner, The bacterial cytoplasm has glass-like properties and is fluidized by metabolic activity, Cell **156**, 183 (2014).

Movie S1: One example of numerical simulations to generate indissoluble biomolecular condensates with elasticity.

Movie S2: Examples of condensates with the bulk moduli below and above their critical values respectively.

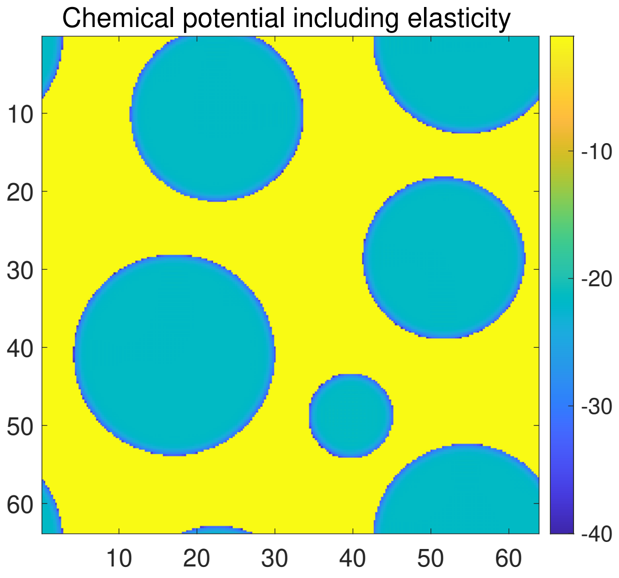


FIG. S1. Heatmap of the effective chemical potential $\mu_{\text{eff}} = f'(\phi) + f'_{\text{el}}(\phi)$, where $f_{\text{el}}(\phi) = -G_B + G_B \ln(\phi_1/\phi)$ is obtained by $\phi f'_{\text{el}}(\phi) - f_{\text{el}}(\phi) = -\sigma_B$. The effective chemical potential is not uniform across the system, which indicates that the elastic energy cannot be simply included in the free energy. In this figure, we take $\phi_0 = 0.45$, $G_B = 20$, $G_S = 20$, and $\phi_c = 0.5$, the same as Figure 2 in the main text.

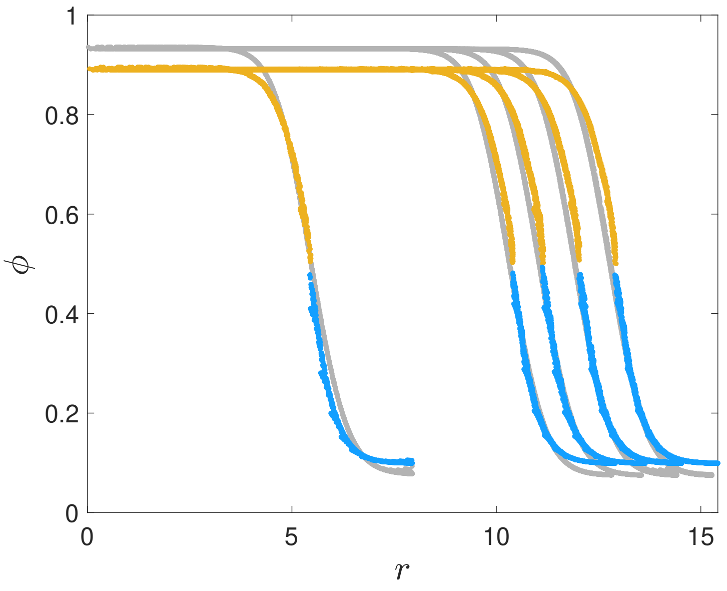


FIG. S2. Numerical verifications of constant radii. The initial $\phi$ are plotted as gray dots and the final $\phi$ are colored dots, which demonstrates that the radii are approximately invariant. This simulation is the same as Figure 2 in the main text.

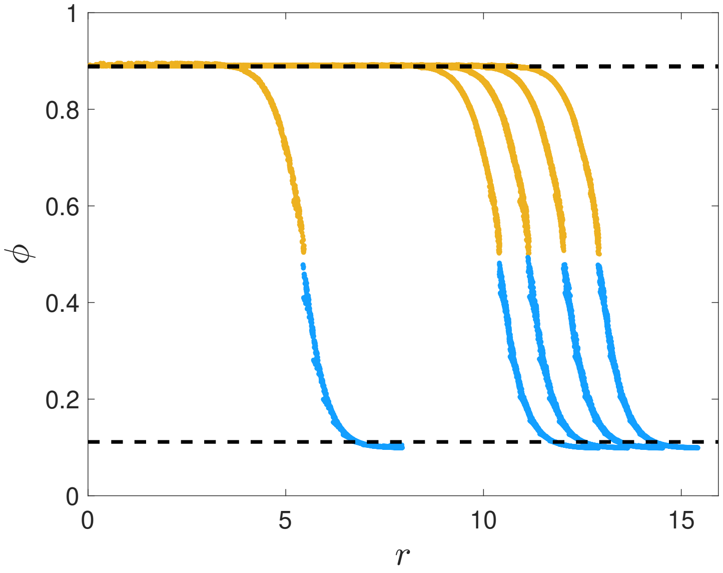


FIG. S3. Theoretical predictions of $\phi$ assuming $\gamma = 0.2$ (black dashed lines), which are very close to the ones in Figure 3b with $\gamma = 0$.

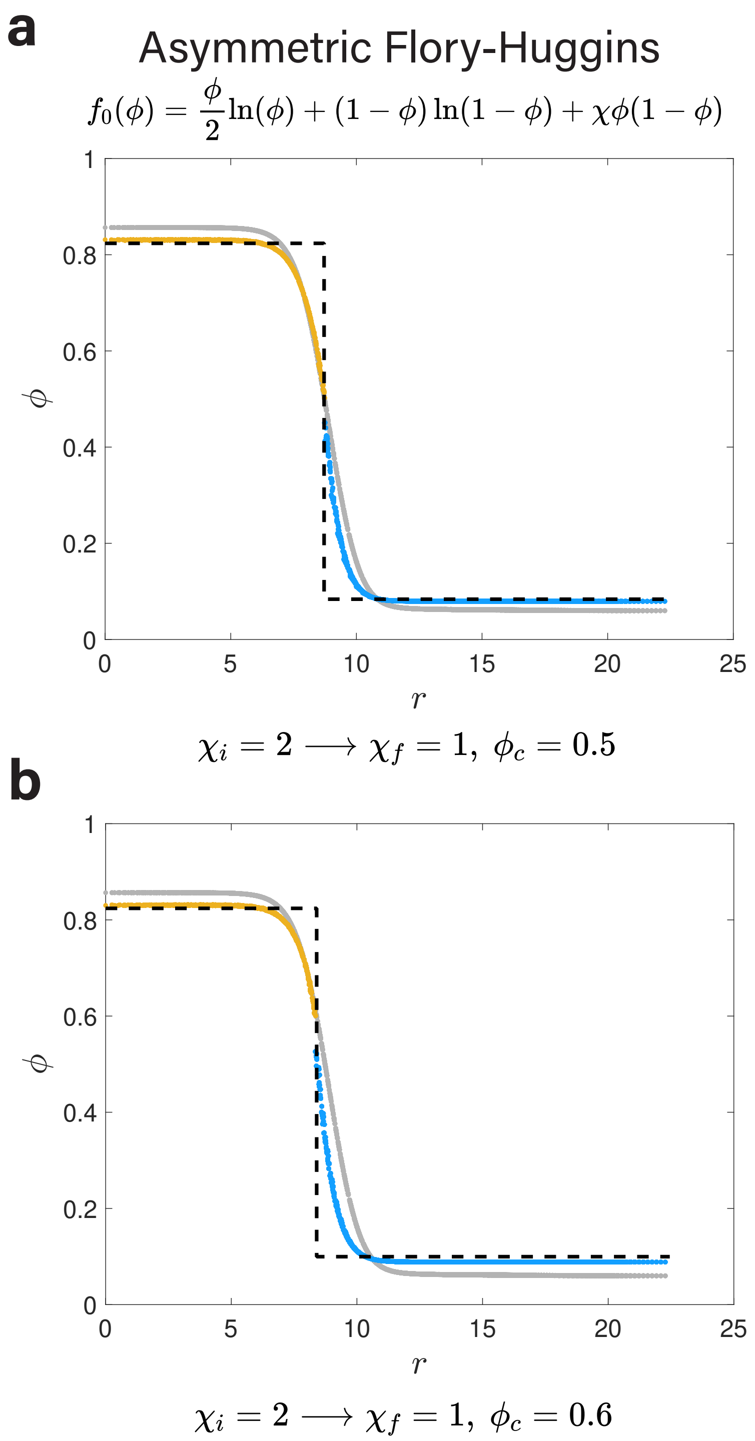


FIG. S4. Simulations using asymmetric free energy. For the polymeric Flory-Huggins free energy with $N = 2$, when the control parameter $\chi$ decreases from 2.0 to 1.0, the initial density field (gray dots) cannot be maintained and the final density field is established (yellow dots above $\phi_c$ and blue dots below $\phi_c$). The black dashed line is the theoretical prediction. In (a) $\phi_c = 0.5$ and in (b) $\phi_c = 0.6$. In both (a) and (b), a single condensate is simulated, and $G_B = 20$, $G_S = 20$, $R_0 = 9$.

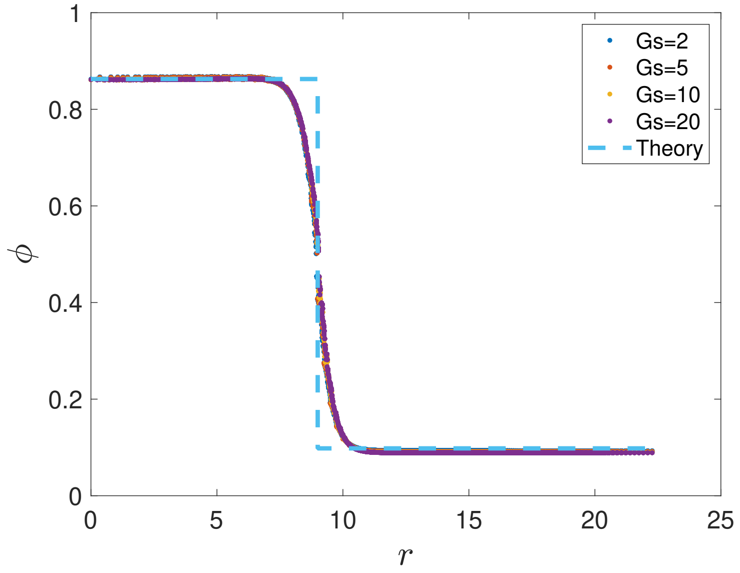


FIG. S5. Simulation results are insensitive to the shear modulus $G_S$. The density fields $\phi$ after decreasing $\chi$ from 3.0 to 1.5 with different $G_S$ are plotted as colored dots, which are almost overlapped showing that the shear modulus rarely affects the simulation results. Here, we take $R_0 = 9$, $G_B = 10$, and $\phi_c = 0.5$.

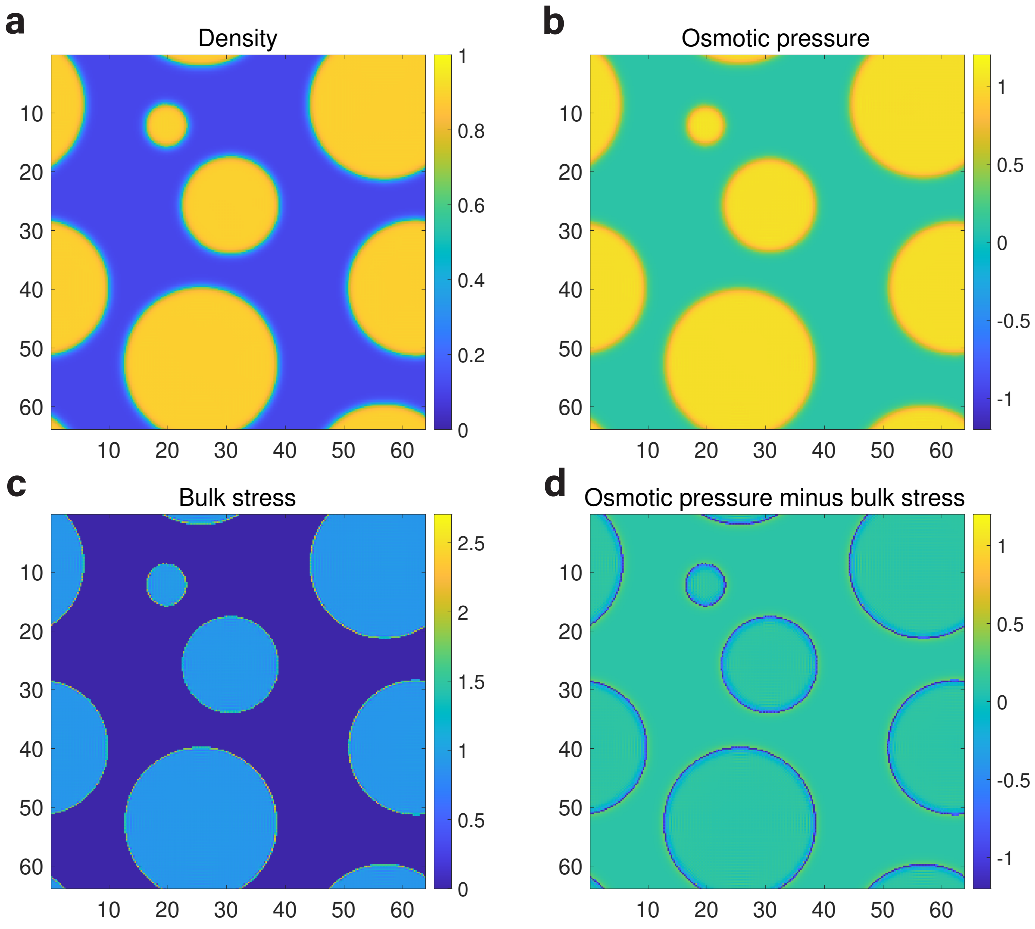


FIG. S6. Simulations of multiple coexisting condensates. (a) The density field $\phi$ after decreasing $\chi$ from 3.0 to 1.5. (b) The osmotic pressure $\Pi$ from the same simulation of (a). (c) The bulk stress $\sigma_B$ from the same simulation of (a). (d) $\Pi - \sigma_B$ from the same simulation of (a). In the simulations of this figure, we take $\phi_0 = 0.45$, $G_B = 20$, $G_S = 20$, and $\phi_c = 0.6$.

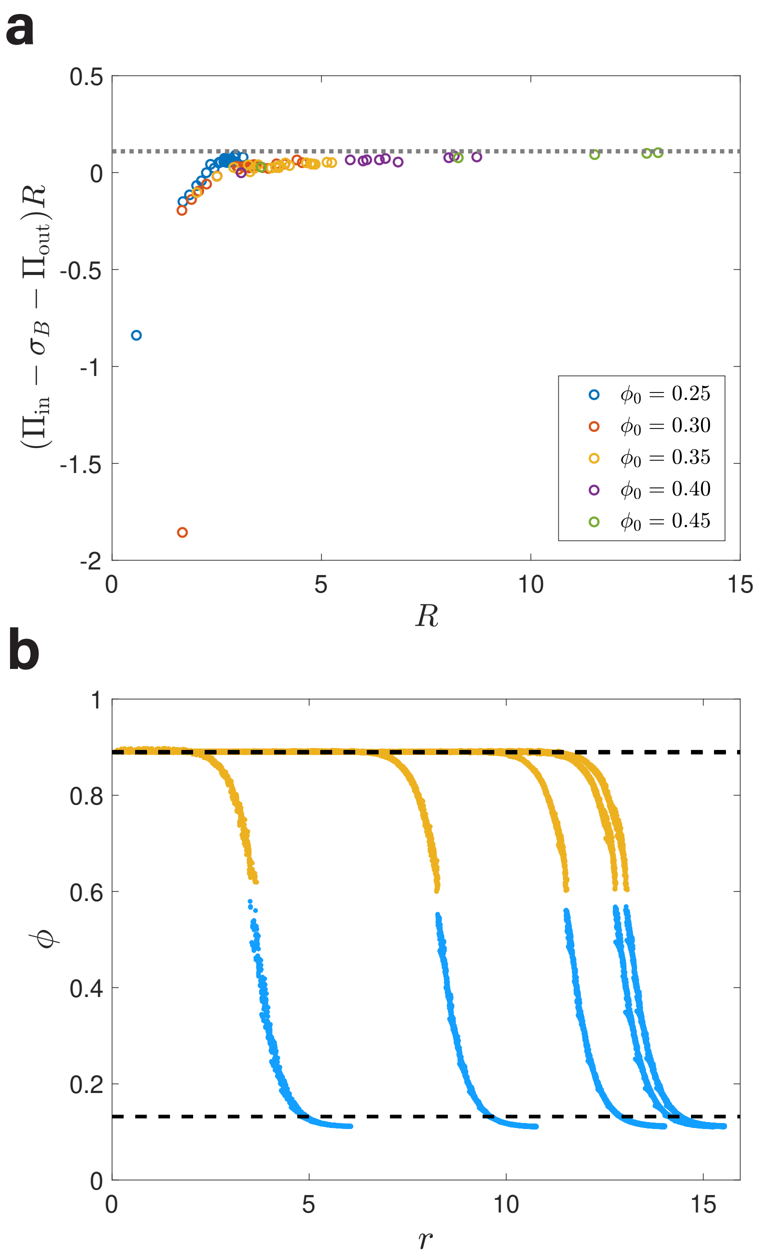


FIG. S7. Computations of the surface tension constant $\gamma$ and predictions of the density field, similar to Figure 3 while $\phi_c = 0.6$. (a) The inferred surface tension constant $\gamma = (\Pi_{\text{in}} - \sigma_B - \Pi_{\text{out}})R$ approaches an asymptotic value in the large radius limit. (b) A comparison of the theoretical predictions of $\phi$ (black dashed lines) and the simulations in Figure S6(a) (yellow dots above $\phi_c$ and blue dots below $\phi_c$). In both (a) and (b), $G_B = 20$, $G_S = 20$, $\phi_c = 0.6$, $\chi_i = 3.0$, $\chi_f = 1.5$. In (b), $\phi_0 = 0.45$.

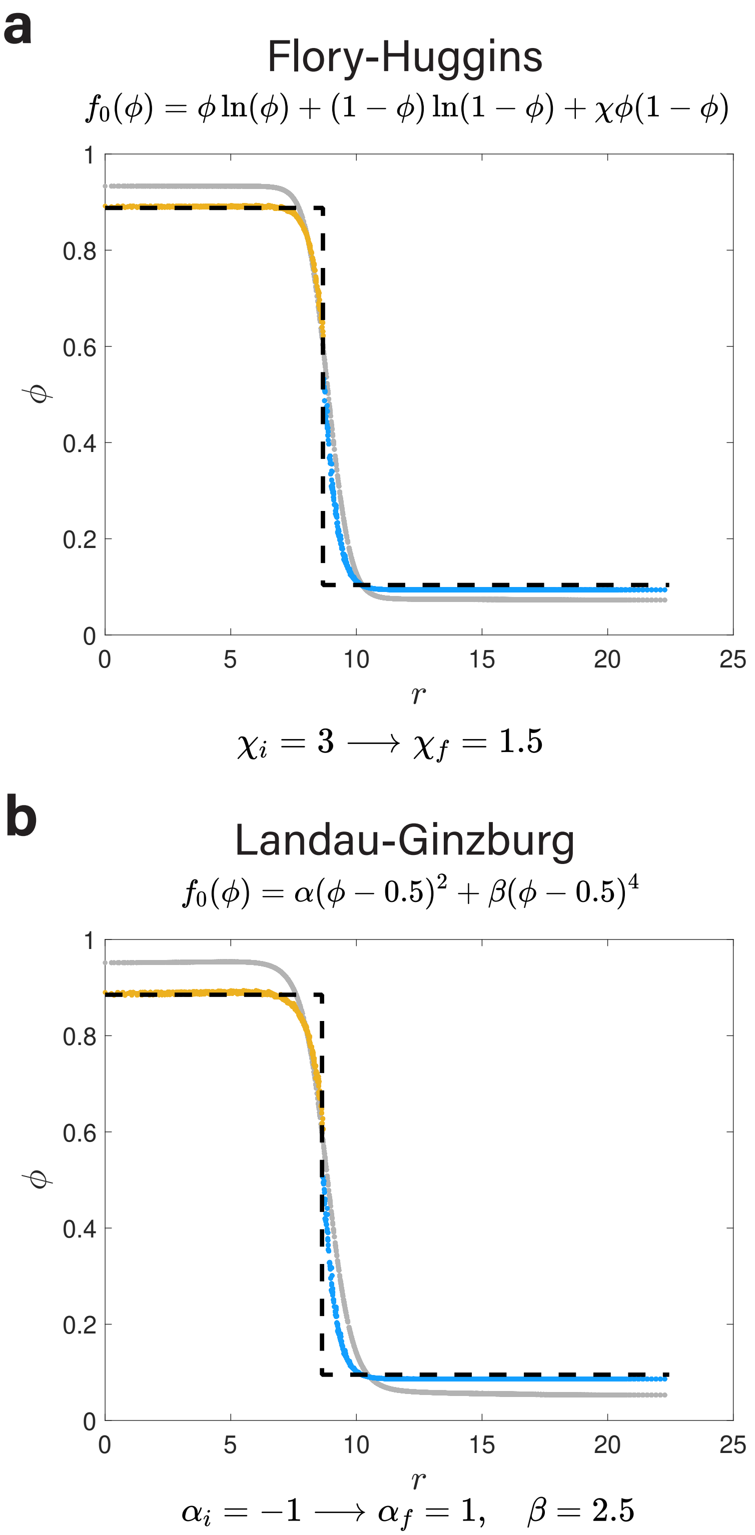


FIG. S8. Simulations using different types of free energy. (a) For the Flory-Huggins free energy, when the control parameter $\chi$ decreases from 3.0 to 1.5, the initial density field (gray dots) cannot be maintained and the final density field is established (yellow dots above $\phi_c$ and blue dots below $\phi_c$). The black dashed line is the theoretical prediction. (b) For the Landau-Ginzburg free energy, the control parameter $\alpha$ increases from $-1$ to 1, and the equilibrium density field can also be predicted by our theories. In both (a) and (b), a single condensate is simulated, and $G_B = 20$, $G_S = 20$, $\phi_c = 0.6$, $R_0 = 9$.

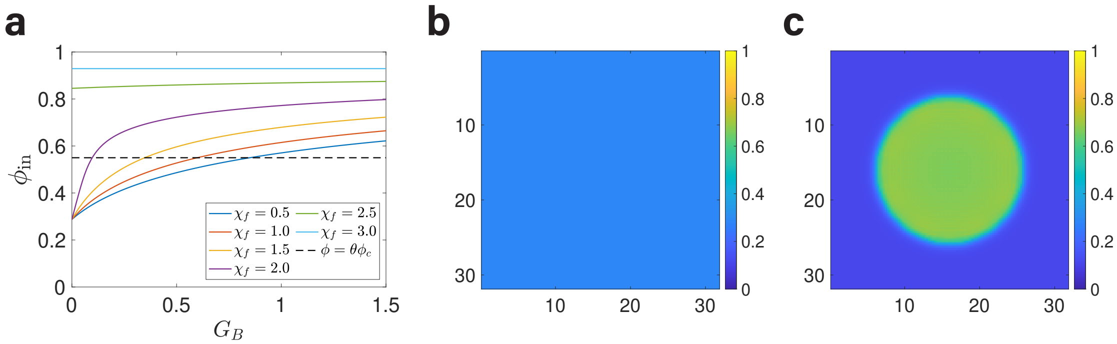


FIG. S9. Theoretical $\phi_{\rm in}$ and examples of condensates under different $G_B$ and $\chi_f$. (a) The theoretical predicted $\phi_{\rm in}$ with $G_B$ under different $\chi_f$. (b) and (c) The density fields $\phi$ for dissolvable and indissoluble cases respectively. In (b), $G_B = 0.4$ and $\chi_f = 0.5$ and the final density is uniform. In (c), $G_B = 1.2$ and $\chi_f = 1.5$ and the condensate is indissoluble due to elasticity.

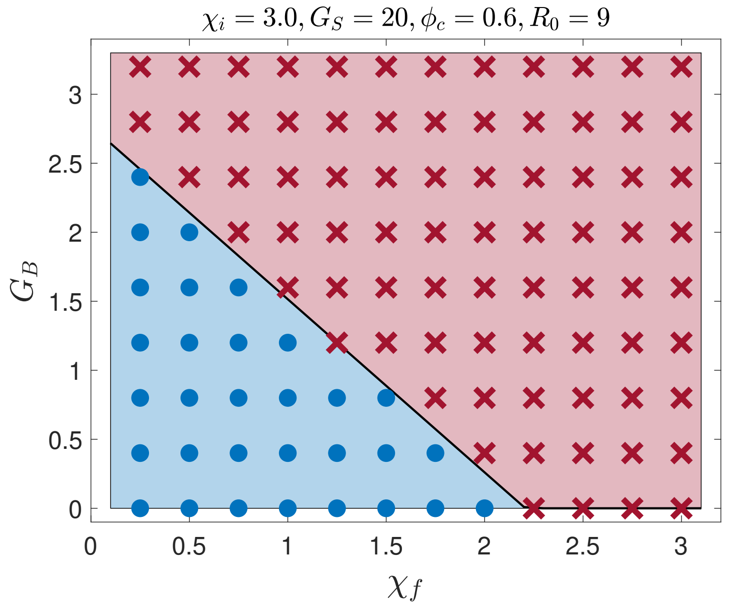


FIG. S10. Phase diagram of condensate stability with control parameters $\chi_f$ and $G_B$. The theoretical predicted $G_{B,c}$ is the black line and the simulation results are the blue dots and red crosses. In this figure, $\phi_c = 0.6$, and the theoretical prediction is obtained with $\theta = 1.1$.

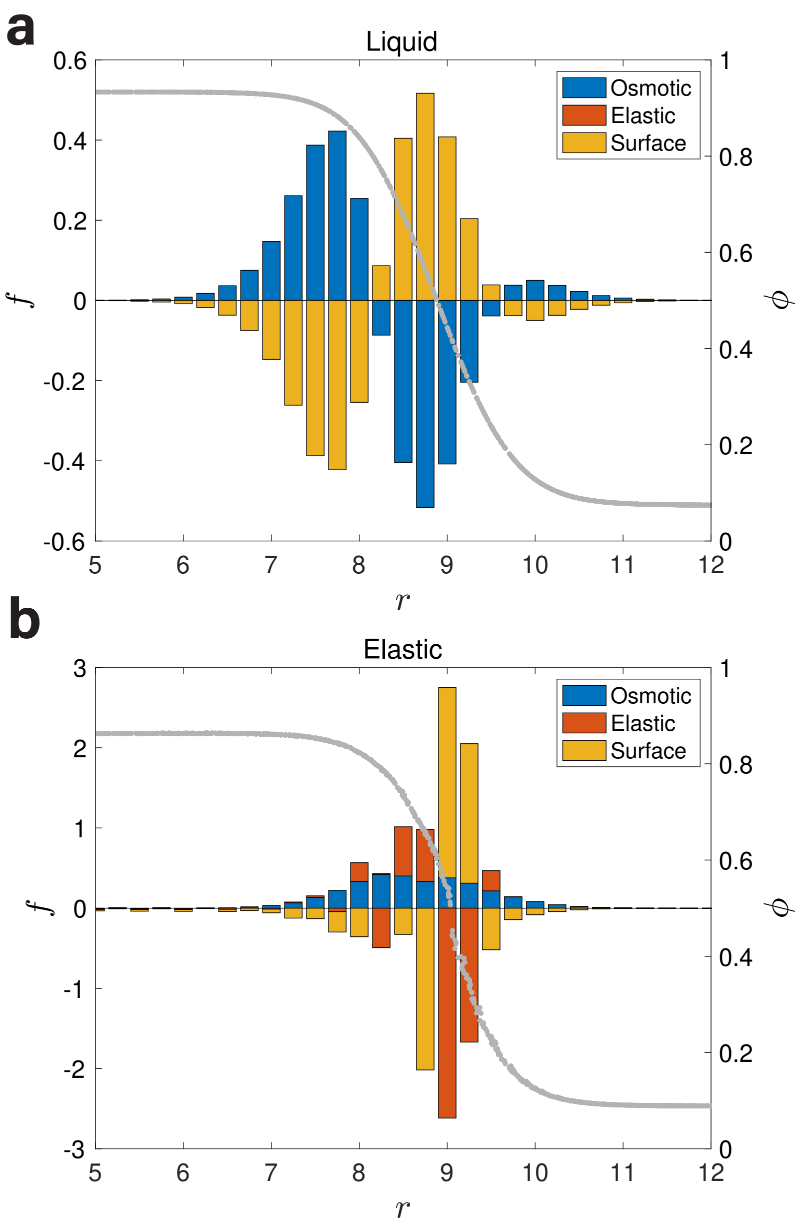


FIG. S11. The force distribution from numerical simulations on the surface of a condensate where a positive (negative) value means an outward (inward) force. (a) The force distributions for a liquid condensate including the osmotic force and the surface tension force, which are separated into three parts. The osmotic force is inward in the second part. (b) The force distributions for an elastic condensate including the osmotic force, the elastic force and the surface tension force, which are roughly separated into three parts. The elastic force is inward in the second part to balance the outward osmotic force. In this simulation, we take $R_0 = 9$, $\chi = 3$ for (a), and $\chi = 1.5$, $G_B = 10$, $G_S = 20$ and $\phi_c = 0.5$ for (b).
